# Some like it hot: adaptation to the urban heat island in common dandelion

**DOI:** 10.1101/2023.06.01.543268

**Authors:** Yannick Woudstra, Ron Kraaiveld, Alger Jorritsma, Kitty Vijverberg, Slavica Ivanovic, Roy Erkens, Heidrun Huber, Barbara Gravendeel, Koen J.F. Verhoeven

**Affiliations:** Netherlands Institute of Ecology, Department of Terrestrial Ecology, Droevendaalsesteeg 10, 6708PB, Wageningen, The Netherlands; Naturalis Biodiversity Center, Evolutionary Ecology, Darwinweg 2, 2333CR, Leiden, The Netherlands; Radboud University Nijmegen, Radboud Institute of Biological and Environmental Sciences, Heyendaalseweg 135, 6500 GL, Nijmegen, The Netherlands; Wageningen University & Research, Laboratory of Genetics, Droevendaalsesteeg 2, 6708PB, Wageningen, The Netherlands; Rijk Zwaan Breeding B.V., Eerste Kruisweg 9, 4793RS Fijnaart, The Netherlands; Maastricht University, Maastricht Science Programme, P.O. Box 616, 6200MD, Maastricht, The Netherlands; Maastricht University, System Earth Science, P.O. Box 616, 6200MD, Maastricht, The Netherlands

**Author notes:** [Contact Information]: Yannick Woudstra, Koen J.F. Verhoeven.

**Keywords:** urban ecology, urbanisation, climate change, anthropogenic environments, human induced rapid evolutionary change (HIREC), rapid selection, disturbed environments, Asteraceae, apomixis

## Abstract

The Urban Heat Island Effect (UHIE) is a globally consistent pressure on species living in cities. Rapid adaptation to the UHIE may be necessary for urban wild flora to persist in cities, but experimental evidence is lacking. Here, we report the first evidence of genetic differentiation in a plant species in response to the UHIE. We collected seeds from common dandelion (*Taraxacum officinale*) individuals along an urban-rural gradient in the city of Amsterdam (The Netherlands). In common-environment greenhouse experiments, we assessed the effect of elevated temperatures on plant growth and the effect of vernalisation treatments on flowering phenology. We found that urban plants accumulate more biomass at higher temperatures and require shorter vernalisation to induce flowering compared to rural plants. Differentiation was also observed between different intra-urban subhabitats, with park plants displaying a higher vernalisation requirement than street plants. Our results show strong differentiation between urban and rural dandelions in temperature-dependent growth and phenology, consistent with adaptive divergence in response to the UHIE. Rapid adaptation to the UHIE may be a potential explanation for the widespread success of dandelions in urban environments.

**Summary statement:** The urban heat island effect (UHIE) is the most prominent and globally consistent characteristic of environmental change due to urbanisation, severely impacting human populations in cities as well as the cohabiting wildlife. Despite the profoundly mitigating effect of vegetation on urban heat, evidence for plant adaptation to the UHIE has been lacking. Here we provide the first experimental evidence to date, demonstrating adaptation in urban dandelions in response to elevated temperatures, similar to the UHIE. We furthermore show an urban-rural differentiation in flowering response to shorter vernalisation times (cold winter period to activate the onset of flowering in early spring). Given the predominantly asexual apomictic mode of reproduction in dandelions, this evolution is likely the result of environmental filtering on a diverse population of clonal genotypes. We conclude that plant adaptation to the UHIE exists and recommend future studies to contrast our findings with those in outcrossing sexual plant systems. Studies of urban heat adaptation can bring impactful contributions to building climate change-resilient environments and plants should be an integral part of this research.

## Introduction

The urban environment is a relatively novel ecosystem that has introduced a significant environmental change to the co-inhabiting and surrounding wild species^1^. Considering the many different stakeholders in urban environments and the rapidity of global urbanisation, it is important to understand the adaptive potential of nature to city life^2^. Eco-evolutionary experiments involving comparisons between rural and urban counterparts can demonstrate if, how and at which rate natural species can integrate into urban environments, important for the safeguarding of ecosystem services^3^. The urban heat island effect (UHIE) is the most well-documented and consistent environmental pressure exerted by urbanisation on nature in and around cities^4^. Adaptation to an urban-rural cline in temperature can therefore be predicted as a hallmark of urban evolution. This has been demonstrated in fungi^5^ and a variety of animal species^6– 11^, providing general evidence for the importance of adaptation to the UHIE. However, adaptation of plants to the UHIE is undemonstrated^12^, despite vegetation being the most important factor in UHIE mitigation. Vegetation provides considerable cooling of urban surface and air temperatures through shading of heat-retaining materials, evapotranspiration, and increased albedo and air circulation^13^. Here, we present the first experimental evidence for a plant species adapting to the UHIE, using common dandelion as a model plant.

The UHIE leads to elevated temperatures, particularly during the growing season, that may be utilised by plants for growth. Temperatures in cities are higher compared to rural areas, caused by the retention and re-radiation of heat from solar radiations by buildings and by the emission of heat from anthropogenic heat sources^14^. The intensity of the UHIE is highly seasonal^15^ and mainly depends on the density of buildings, intensity of traffic, presence of industrial activity and the mitigation of heat by water and vegetation^4^. For example, in the capital of the Netherlands, Amsterdam, the UHIE leads to an annual average temperature increase of >2 °C in the city centre (Figure 1A), while a hot summer day can elevate this difference to up to 9 °C for ambient temperatures and up to 20 °C for surface temperatures^16^. These circumstances would selectively favour species^7^ and genotypes that are optimised for growth at elevated temperatures.

**Fig. 1:**
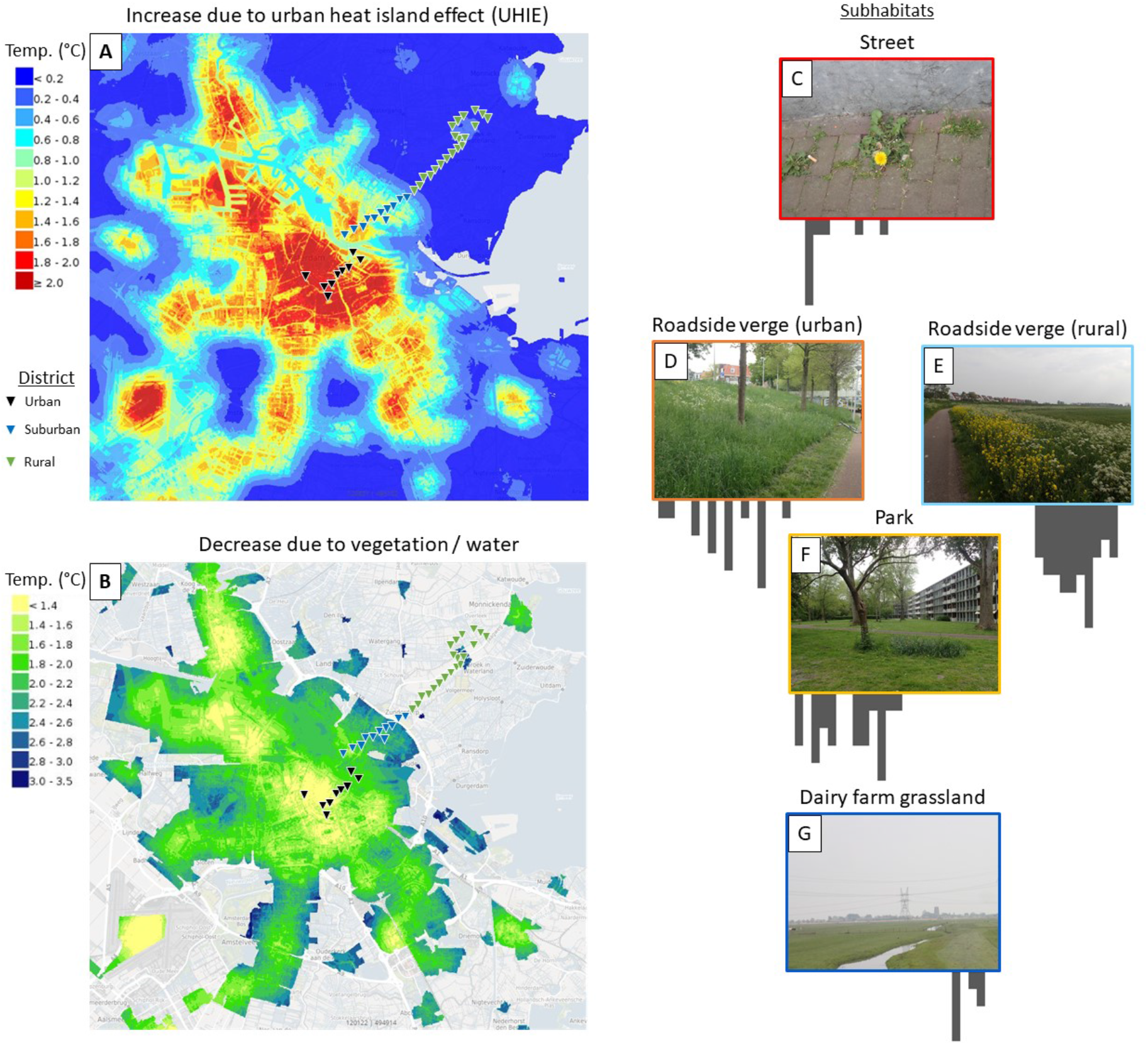
Overview of the urban heat island effect (UHIE) in Amsterdam, The Netherlands. Locations in the sampling transect are indicated with small black triangles and are coloured based on the district they occur in. Two aspects of the urban heat island are shown: **A** - net increase in temperature due to the UHIE (retention of heat by buildings, streets and other man-made impermeable surfaces); **B** - effect of UHIE mitigation by surrounding vegetation and water. Both maps indicate annual averages. Mitigating effects shown in **B** are already discounted in the temperature increases shown in **A**. The transect was further divided into subhabitats based on field observations of the immediate surroundings (**C-G**). Histograms under the subhabitat pictures reflect the distribution of each subhabitat along the transect ranging from the start (left side of each picture) to the end (right side of each picture)) of the transect, based on 120 individual accessions. Subhabitat definitions are detailed in Materials & Methods section 1. Source for maps: Rijksinstituut voor Volksgezondheid & Milieu^29^, https://www.atlasleefomgeving.nl/stedelijk-hitte-eiland-effect-uhi-1.

The UHIE can also affect the timing and duration of the growth season^17^, creating differential phenological optima between urban and rural habitats. In cities in the Northern hemisphere, winter surface temperatures are on average 1-2 °C higher and warm up earlier in the year compared to rural sites^15^. Dandelions in north-western Europe are mostly apomictic; despite their asexual nature, dandelions require flowering for reproduction as the clonal (apomictic) seeds are formed by parthenogenesis from unreduced egg cells^18^. An important determinant of dandelion flowering phenology is through vernalisation: extended exposure to low temperatures is required to enable the transition to flowering. Populations of clonal dandelions display large genotypic variation in flowering phenology^19,20^, providing a genetic reservoir for phenological adaptation. Differential selection on vernalisation requirements and flowering time can therefore be expected along urban-rural clines in relation to the UHIE.

The common dandelion (*Taraxacum officinale* F.H.Wigg.) is one of the most widespread plant species in temperate areas and occurs prolifically both in rural areas around cities and within cities, where it functions as one of the most important food sources for insects^21^. Nearly all common dandelions in Northern Europe are triploid apomictic plants^22^ and populations are highly diverse assemblages of different clonal genotypes^23^ upon which natural selection can act. Such diverse clonal assemblages can show highly repeatable^24^ and rapid adaptive responses to selection^25^, producing population-level adaptation by environmental filtering of (clonal) genotypes^26,27^. Apomixis also allows for easy replication of genotypes for phenotypic evaluations in common garden experiments, to test whether observed phenotypic divergence between urban and rural plants has a genetic basis. The above reasons make the common dandelion an ideal model for experimental studies on urban adaptation in plants.

To fill the knowledge gap on plant adaptation to the UHIE, we experimentally tested temperature-related adaptation in common dandelions along an urban-rural transect. Specifically, we evaluated growth response to elevated temperatures and flowering response to different lengths of vernalisation period. We did this in controlled common garden experiments using clonal seeds collected along an urban-rural transect in the city of Amsterdam, The Netherlands. In order to discover the spatial scale at which adaptation occurs along this urban-rural cline, we analysed phenotypic divergence between plants in relation to (sub)urban versus rural areas, subhabitats within the urban-rural mosaic (street, urban roadside verge, park, rural roadside verge and dairy farm grassland), and distance to the centre. We hypothesised that urban plants grow better at elevated temperatures and require shorter vernalisation periods compared to rural plants. We show that rapid adaptation to the UHIE occurs in dandelions, indicating that natural selection on diverse clonal assemblages can facilitate plant resilience to urban heat.

## Materials & Methods

### 1. Amsterdam urban-rural transect

We sampled common dandelion plants (*Taraxacum officinale* F.H.Wigg.) along an urban-rural gradient in Amsterdam, The Netherlands, following the protocol from the GLobal Urban Evolution project (GLUE, https://www.globalurbanevolution.com/). Briefly, GLUE transects were designed for sampling white clover (*Trifolium repens* L.) and are defined as 1) covering a continuous urbanisation gradient, 2) containing an equal amount of distance and number of sampling locations within and outside the urban area, and 3) contain a minimum total of 40 sampling locations with a minimum distance of 200m between sampling locations. Due to the natural habitat of white clover, a high proportion of sampling locations occur within urban green spaces (parks and mowed lawns). The Amsterdam transect (Figure 1) has 40 sampling locations and runs from the approximate geographical centre of the city (sampling location 1: Sarphati Park, Oud-Zuid) through suburban areas (Amsterdam-Noord) to a small rural village (sampling location 40: Broek in Waterland), measuring a total distance of 12.03 km. In the late spring season (22-26 April 2020) we collected dandelion seed heads from three plants at each sampling location, aiming for a minimum of 50 m distance between individuals. In total, seed heads were collected from 120 accessions (40 sampling locations x 3 individuals per location).

To study differentiation among urban and rural dandelions at different spatial scales, we annotated the accessions collected along the transect in three different ways. Firstly, we divided the transect in three general districts (Figure 1A-B) comprising urban (locations 1-9), suburban (locations 10-20) and rural (locations 21-40) districts. Secondly, we defined subhabitats (Figure 1C-G) occurring along the urban-rural transect based on our own observations during the dandelion flowering season: I) street/pavement - a continuous impermeable surface of man-made structure (i.e., a road); II) urban roadside verge - a green strip (planted border or lawn) next to an urban road; III) city park - a continuous green zone within the (sub)urban perimeter where car traffic is excluded; IV) rural roadside verge - a green strip of (naturalised) vegetation along rural road (or bicycle lane); V) dairy farm grassland - continuous zone of grassland vegetation (usually a monoculture of English ryegrass (*Lolium perenne* L.) with a few wildflowers) that is grazed by dairy cattle or frequently mown for hay harvesting and typically highly fertilised. Finally, we calculated the distance for each accession to the start of the transect (sampling location 1) as an approximation of distance from the centre of the heat island. We recorded the GPS locations in the field during collection using a smartphone (Motorola Moto G5, Motorola Inc., Chicago, IL) and measured distance in a straight line to the start of the transect using an online calculator (https://www.gps-coordinates.net/distance). An overview of categorical divisions for all accessions can be found in Suppl. Mat. I.

Several taxonomic sections of *Taraxacum* co-exist in the geographic research area that are not always easily distinguishable in the field. We aimed to restrict our analysis as much as possible to one section: *Taraxacum* F.H.Wigg. sect. *Taraxacum* (formerly known as *T*. sect. *Ruderalia* Kirschner et al.), as this was taxonomically determined to be the section of “agamospermous polyploid microspecies and the closely related sexual diploids generally known as common dandelions”^28^. We used the position of outer involucral bracts in the mature inflorescence morphology as a diagnostic character, marking those with dispersed or strongly reflexed bracts as *T*. sect. *Taraxacum*. This selection was performed with the inflorescences obtained in the long vernalisation experiment as nearly all plants reached flowering here. Ten accessions were thereby omitted from further analysis.

### 2. Vernalisation experiments

To determine differential vernalisation requirements between urban and rural dandelions we subjected all accessions from the transect to three different lengths of a continuous cold period (4°C): a long vernalisation period of 109 days, a short period of 36 days and an absence of vernalisation (control). We considered not flowering as a lack of sufficient vernalisation, considering the long vernalisation requirement in wild dandelions in general^30,31^.

#### a. Long vernalisation

Seeds collected in the field were germinated on agar plates for 11 days (20 °C, 16h light / 16 °C 8h dark) in ECD01 incubators (Snijders Labs, Tilburg, The Netherlands). For each accession, one seedling was transplanted into a 9.3×9.3×10 cm pot containing, volume wise, 50% potting soil and 50% fine sand. Seedlings were then propagated in the greenhouse for 28 days and watered once or twice per week, as required. Subsequently, all plants were given a vernalisation treatment by putting them in a cold chamber for 109 days (4 °C, 16h light / 8h dark), based on standardised vernalisation protocols for dandelions^31^. Plants were then placed back into the greenhouse in random order, with watering every 2-3 days. Once a week we supplied a 2x diluted Hoagland nutrient solution^32^ for optimal growth. Flowering time was recorded in days from the time of re-entering the greenhouse (day 0) until the emergence of the first yellow petals from the first inflorescence. Flowering was monitored for 107 days after 1 February 2021; plants that had not flowered at that moment were scored as non-flowering.

#### b. Short versus no vernalisation

In a second vernalisation experiment a short cold treatment (36 days: 4 °C, 16h light / 8h dark) was applied. Field collected seeds were newly germinated (two per accession) and propagated from seedlings to small plants, using the conditions described in section 2a, with the exception that a substrate mixture of 80% potting soil and 20% pumice was used. One batch of plants was subjected to the short vernalisation treatment (36 days), while the other batch remained in the greenhouse to serve as a control for absence of vernalisation. Plants were placed on greenhouse benches in random order within treatments. Water and fertiliser supply followed the protocol described in section 2a, supplemented by a singular supply of 1.3 grams Osmocote® Exact Mini 5-6M slow-release fertilisation pellets (ICL Group Ltd, Tel-Aviv, Israel) per pot. Flowering time was recorded in days from the day of (re-)entering the greenhouse (day 0). Flowering was monitored for 118 days after 16 March 2021; plants that had not flowered at that moment were scored as non-flowering.

### 3. Heat experiment

For each flowering plant in the long vernalisation experiment (Section 2a), the first developing inflorescence was used to produce clonal seeds for the heat experiment. Any potential cross-pollination was prevented by putting small paper bags over the inflorescences as soon as they opened. Bags were removed after 3-4 days when seeds started to develop. This can be seen as confirmation of apomixis, considering that common dandelions are self-incompatible^33^, and therefore self-pollination can be discounted as a potential mode of fertilisation.

To test whether urban and rural dandelions differ in their growth response to elevated temperatures, we measured seedling biomass accumulation in urban and rural dandelions at a range of different temperatures. For this experiment, we utilised clonal seeds harvested from the first seed heads of plants grown in the common environment of the long vernalisation experiment (section 2a). By using seeds that were generated in a common greenhouse environment, we minimise environmental maternal effects on seed quality as a source of phenotypic variation in experimental plants. Seed germination and seedling propagation were performed using the conditions described in section 2a, with the exception that smaller pots (7×7×8 cm) and a shorter seedling propagation time (7 days) were used. The length of the 3^rd^ leaf was measured as an estimator of plant size at the beginning of the temperature treatment. Using MC 1750VHO climate cabinets (Snijders Labs), we grew the seedlings at 20, 26, 32 and 38 °C for 11 days (60% humidity, 16h light / 8h dark). For each temperature, we used three replicate blocks with one plant from each accession placed in randomised order, leading to a total of three replicates for each accession. Because only two climate cabinets were available at one time, the replicates were split up in time slots. First, two temperature treatments were done for one replicate, whereafter the other two temperature treatments were performed. The second and third replicate blocks were then performed in the same manner. For this reason, seed germination and seedling propagation was performed in batches, to ensure equal germination propagation time for all plants in all temperature treatments. Plants were harvested after the temperature treatment, the roots were washed, and the whole plant was dried (70 °C, 1 week) before weighing the dry biomass.

### 4. Statistical analysis

We used three different analysis approaches to test if the probability of flowering differed between plants according to their collection position along the transect. First, we divided the transect in 25 windows of 500m, and we used logistic regressions to test if the proportion of plants that flowered in our vernalization experiments within a 500m window showed a gradual change along the transect. Logistic regressions were performed using PROC GENMOD (SAS OnDemand for Academics, SAS Institute Inc., Cary, NC) using Wald Chi-square tests. Second, we used Fisher’s exact tests (PROC FREQ, SAS OnDemand for Academics) to test if the proportion of plants that flowered in our vernalization experiments differed between the three main transect districts: urban, suburban, and rural. Third, we used Fisher’s exact tests to test if the proportion of plants that flowered in our vernalization experiments differed between the five subhabitats: street, urban roadside verge, park, rural roadside verge and dairy farm grassland. For all analyses, we fitted separate models to data of the three separate vernalization treatments (no vernalization, short vernalization and long vernalization).

For the heat experiment, we used mixed linear models to test the effects of temperature, collection position, and their interaction, on total plant biomass. In all models we included replicate block and temperature treatment as categorical fixed factors, 3^rd^ leaf length at the start of the temperature treatment as a continuous fixed cofactor, and accession identity as a random factor. In all models we included ‘collection location of the accession’ and its interaction with temperature treatment as fixed factors, where three separate models used different definitions of ‘collection location’: (1) distance from the start position of the transect (continuous cofactor); (2) main transect district (urban, suburban or rural; categorical factor); and (3) subhabitat type (street, urban roadside verge, park, rural roadside verge and dairy farm grassland; categorical factor). Mixed models were performed using PROC MIXED (SAS OnDemand for Academics, SAS Institute Inc., Cary, NC). After plotting temperature treatment results as a function of distance to the start of the transect, linear regressions were fitted in R ^34^ using the function “stat_poly_line” from the R package “ggpmisc”. All p-values reported in this manuscript correspond to two-sided statistical testing.

### 5. Data availability

All experimental data is available as a supplement to this publication (Suppl. Mat. II).

### 6. Code availability

All code used for statistical analysis is available as a supplement to this publication (Suppl. Mat. III).

## Results

We used seeds from dandelions collected along an urban-rural transect in and around Amsterdam to analyse two traits in which we expected to find evidence of adaptation to the urban heat island: vernalisation requirement (length of cold period necessary to induce flowering) and growth response to elevated temperatures (biomass accumulation). A total of 120 accessions were collected and grown in a common garden environment. We restricted our analyses to sect. *Taraxacum*, thereby omitting 10 accessions based on deviating flowering morphology (Suppl. Mat. I for details).

### 1. Flowering response to shortened vernalisation periods

Rural plants had a higher vernalisation requirement than urban plants (Fig. 2). Only 18 plants flowered in absence of a vernalisation treatment, irrespective of sampling location of the accessions (Fig. 2A, D, G). However, urban-rural differences were expressed under vernalisation treatments. Specifically, a short vernalization treatment was sufficient to induce flowering in the majority of the urban plants but not of the rural plants (Fig 2H). All urban plants except one flowered after long vernalisation (Fig. 2C), whereas a quarter of the rural plants still did not reach flowering (P=0.031, Fig. 2I). The probability of flowering under this treatment steadily decreased with distance to the centre of the UHI but dropped significantly faster in the rural area (P=0.015, dotted line in Fig. 2C). We conclude that our vernalization treatments were long enough to satisfy the vernalization requirement for nearly all urban accessions, but not for all rural accessions. For plants that did flower, no significant flowering time differences were observed between urban and rural plants when vernalisation was applied (Suppl. Mat. IV). Urban plants flowered slightly later than rural plants when no vernalisation was applied. This was mainly the effect of earlier flowering in rural roadside verge plants in the absence of vernalisation.

**Fig. 2:**
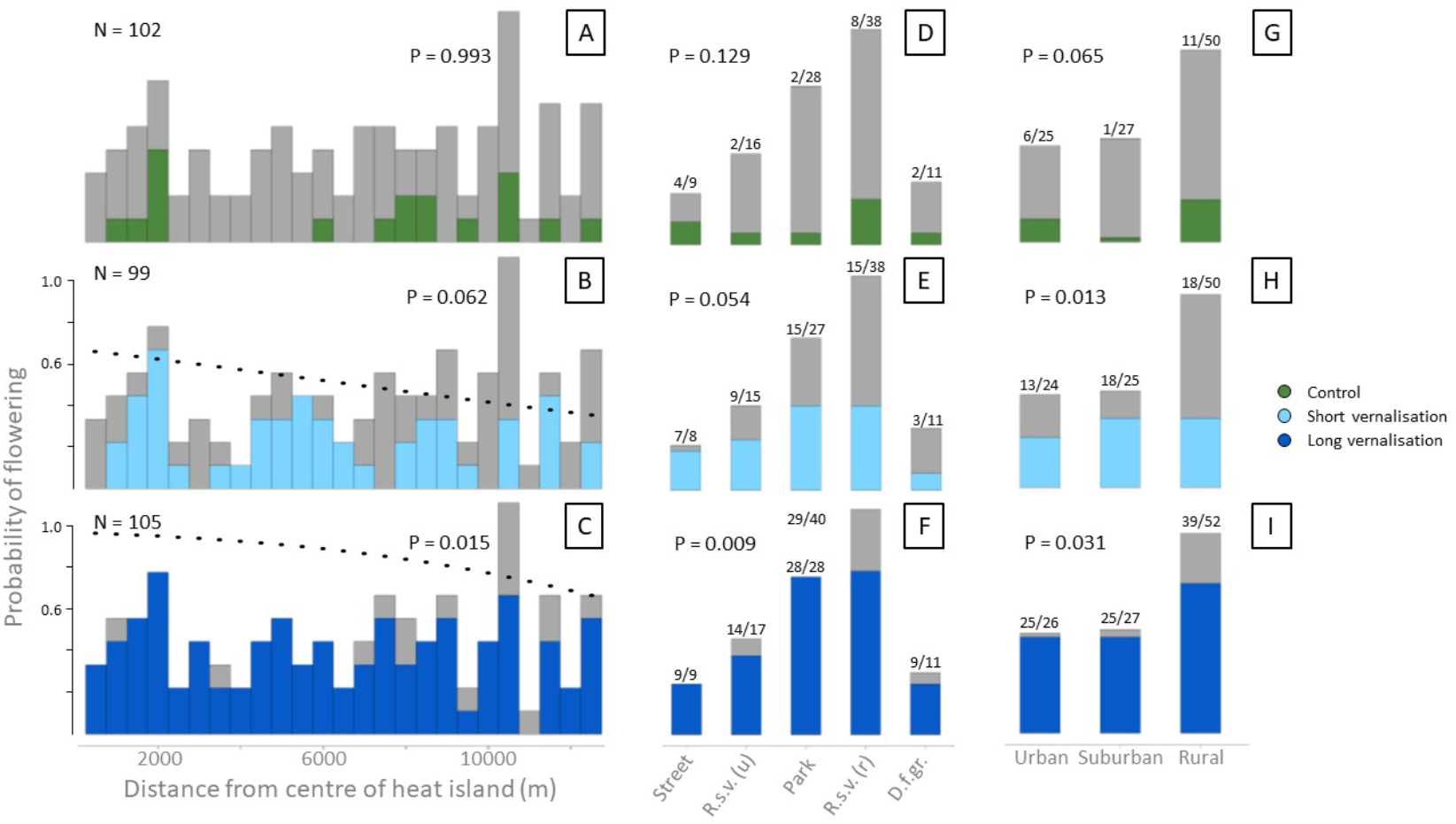
Proportions of dandelions flowering under different vernalisation treatments, for plants originating from different locations along the urban-rural sampling transect. The number of flowering plants is plotted in colour against the total number of plants in the experiment (grey) in relation to: distance from the centre of the heat island (defined as the starting point of the transect) (A-C) with a bin size of 500m; urban, suburban and rural districts (D-F); and subhabitats (G-I). Top panels (A, D, G): no vernalization; middle panels (B, E, H): short (36 days) vernalization treatment; bottom panels (C, F, I): long (109 days) vernalization treatment. For panels B and C, probabilities of flowering at each distance from the centre of the heat island are plotted as predicted by the logistic regression models. Probability values indicate significance of flowering probability differences along the transect (A-C) or between habitats or districts (D-I).

We observed a significantly different response to vernalisation between subhabitats, where a >7-fold increase was observed in flowering city park plants (Fig. 2E). The response was less strong among street plants (from 4/9 to 7/8 to 9/9) compared to plants found in urban roadside verges (from 2/16 to 9/15 to 14/17) and city parks (from 2/28 to 15/27 to 28/28). The response was lowest in dairy farm grassland plants where only one additional plant was found flowering after a short vernalisation treatment. Streets and city parks were the only subhabitats where every plant reached flowering in the greenhouse. The proportion of non-flowering plants was highest among rural roadside verges, with 11 out of 40 plants failing to flower even after a long vernalisation treatment.

### 2. Growth response to elevated temperatures

The plant growth response to elevated temperatures differed along the transect (Fig. 3A). Increasing the growing temperature from 20 °C to 26 °C and 32 °C caused a significantly stronger growth increase in accessions collected close to the centre of the UHI compared to accessions collected outside the city. This resulted in a higher biomass for urban plants (∼11g more than rural plants) at 26 °C, while rural plants accumulated slightly more (∼2 g) biomass than urban plants at 20 °C (Fig. 3C). Differences disappeared when increasing to 38 °C, where sub-optimal growing conditions were reached for all accessions.

**Fig. 3:**
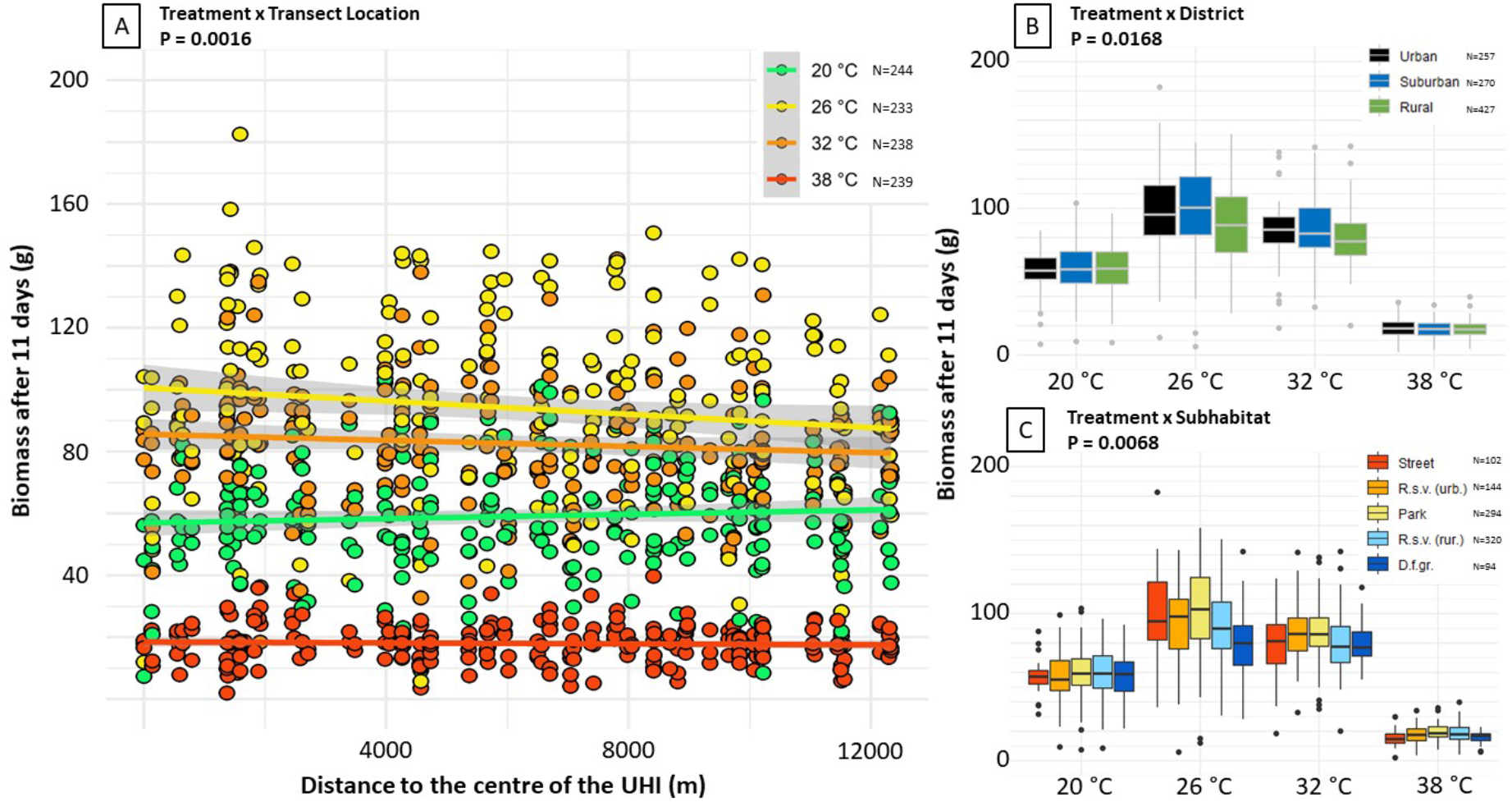
Growth of dandelion seedlings at different elevated temperatures. All accessions collected along the Amsterdam transect were subjected to three elevated temperatures, and a control temperature (20 °C). Accumulated biomass was measured after 11 days as the dry weight of the plants and is visualised in relation to three spatial scales: **A** - distance to the centre of the urban heat island (UHI), with coloured lines indicating the linear regression for biomass at the indicated temperature and grey shading around the lines indicating the standard error; **B** - urban, suburban and rural districts; **C** - subhabitats defined within the transect (Fig. 1C-G), where ‘R.s.v.’ abbreviates ‘roadside verge’ (urban and rural) and ‘D.f.gr.’ abbreviates ‘dairy farm grassland’. Definitions of the different variables are detailed in Materials & Methods section 1.

Differential growth responses to elevated temperatures between rural and urban plants were also visible at the other spatial scales we analysed. Urban and suburban plants showed a higher growth promotion with temperature increases from 20 to 26 °C and 32 °C than rural plants did (Fig. 3B). Similarly, growth differences between subhabitats were negligible at 20 °C but became apparent at higher temperatures, especially at 26 °C, where plants from urban habitats attained higher biomass than plants from rural subhabitats (Fig. 3C). Accessions from rural dairy farm grasslands profited least from the temperature increase to 26 °C.

## Discussion

The Urban Heat Island Effect (UHIE) is the globally most consistent consequence of urbanisation^14^, considerably affecting human health^35,36^ as well as the co-inhabiting biodiversity^1^. Given the high rate of global urbanisation^37^ and the mitigating effect that plants have on the UHIE^13^, it is relevant to understand the adaptive capacity of wild plant species to this novel environment. In this study we presented the first experimental evidence for plant adaptation to the UHIE. We demonstrated adaptive divergence in growth response to increased temperatures using common garden experiments with dandelions from the city of Amsterdam, The Netherlands. Urban plants responded more strongly to increases in temperature than rural plants, by showing a larger increase in biomass accumulation (Fig. 3). In addition, we found differential vernalisation requirements along the urban-rural gradient, where urban and suburban plants generally required less vernalisation than rural plants. The UHIE leads to higher temperatures, especially in summer^15^, during the growing season of plants^17^ and shorter vernalisation exposure due to milder winters^14^. Our results are therefore direct evidence of genetic differentiation in the urban-rural population due to the UHIE. Our temperature experiment furthermore shows a fitness increase (improved biomass accumulation), demonstrating this differentiation as adaptive.

While urban plant adaptation has been demonstrated in other systems^12^, evidence for plant adaptation to urban heat has remained elusive. The characteristics of cities that are most impactful on vegetation are increased temperatures, impervious surface cover, habitat fragmentation, changes in herbivore communities, and light, air and soil pollution. Habitat fragmentation was found to alter dispersal characteristics in urban plants of holy hawksbeard (*Crepis sancta*)^38^ and urban-rural adaptation to changes in herbivory pressure has been demonstrated in white clover (*Trifolium repens*)^39^ as well as dandelions^40^. More generally speaking, the combined pressures of the urban environment (pan-urban effects) have led to phenological adaptation in common ragweed (*Ambrosia artemisiifolia*)^41^ and changes in life-history traits in *Arabidopsis thaliana*^27^. However, none of the above studies have directly explored the effects of urban heat on plant adaptation, making our work a unique milestone contribution to the field of urban plant adaptation.

Proof of adaptation requires the demonstration of genetic differentiation as well as fitness gain^12^, meaning that divergence alone is not enough to claim adaptive evolution. We interpret our results as evidence for adaptation in growth response to urban heat and potential adaptation in flowering phenology to shorter vernalisation exposure in mild urban winters. Firstly, by using common garden experimental designs for phenotypic analysis, genetic differentiation is demonstrated for both traits. Due to the apomictic clonal mode of reproduction in dandelions, the greenhouse-grown plants are genetically identical to the wild mother plant from which the seeds were collected. Conclusions from such common garden experiments are often complicated by the presence of maternal effects^42^, which are phenotypic effects in offspring that reflect the environmental conditions in which the maternal parent was growing, e.g. by means of seed quality, and not reflecting the offspring’s genotype. Differences between urban and rural plants grown in a common environment may therefore (partly) reflect differences between mother plants and their respective environmental plasticity^42^. Such effects can be discounted in the results from our heat response results as the experiment was conducted using clonal second generation seeds harvested from greenhouse-grown plants, strengthening our evidence for adaptive evolution. Secondly, a fitness gain can be inferred from the heat treatment results, as urban plants generated more biomass than rural plants in response to temperature increases. The ability to generate more biomass is strongly linked with higher fecundity in plants, making it a suitable proxy for fitness in adaptation studies^43^. Higher biomass production in urban plants was temperature-dependent, and not consistently observed at all temperatures, thus strengthening the conclusion that this adaptation is a response to the UHIE.

We interpret the lower vernalisation requirement in urban plants (Fig. 2), on the other hand, as tentative evidence for adaptation. Previous studies have demonstrated shifts in flowering phenology for the overall urban plant community in relation to the UHIE^44–46^ but have always relied on field observations in whole ecosystems. Such observations cannot distinguish between phenotypic plasticity and genetic divergence. Our study overcomes these limitations by demonstrating a differential response to shorter vernalisation periods in a single plant species using a common environment. Overall, the urban dandelion population displays a lower vernalisation requirement than the rural population, which is an expected result of selection in response to shorter and milder urban winters due to the UHIE. Such adaptation could result from the selective loss of genotypes with long vernalization requirements, which might not be satisfied in cities. Our results on vernalisation requirement differences between intra-urban and -rural subhabitats emphasise the complexity of the urban-rural mosaic, including the UHIE. Large city parks, for example, present relatively “cool” islands within the urban heat island (Fig. 1A), which is reflected in a higher vernalisation requirement in these plants compared to urban street plants (Fig. 2D-F). While our vernalisation results are consistent with the hypothesis of adaptation to the UHIE, additional empirical evidence on the fitness benefits of shorter vernalisation requirements in cities is desired to strengthen the adaptive interpretation of the observed genetic divergence in this trait.

Having demonstrated rapid adaptation to urban heat in apomictic dandelions, a relevant question is to what extent our results are generalizable to other plant species. We used the clonal mode of reproduction found in dandelions to our advantage here as it allowed us to establish a common environment experiment with genetically identical copies of the wild plants. Additionally, rapid adaptation may be facilitated in apomictic dandelion populations as they form highly diverse clonal assemblages^23^. Such systems can respond rapidly to selection by environmental filtering of clones^25^, however, it is unclear if the long-term populations’ adaptive potential remains sufficient in these asexual assemblages. It would therefore be of interest to study urban heat adaptation in an outcrossing sexual species, for contrast with the clonal assemblages studied here. Clonality and asexual reproduction are predicted as favourable traits in urban environments due to the lower abundance and diversity of pollinators^47,48^ and the lower availability of nearby sexual mates. Diverse clonal assemblages such as dandelions may therefore have a competitive advantage over sexually reproducing plants.

Exotic plant species are characteristically abundant in the urban habitat, either due to their thriving in disturbed habitats^47^ or their use as ornamentals in gardens and parks in temperate cities, providing opportunities for dispersal as garden escapees^49^. Exotic plants are a concern as they initially tend to decrease support for native insect biodiversity^50,51^. Naturally occurring species on the other hand, such as dandelions in Europe, provide an important food source for pollen- and nectar-feeding insects in urban meadows^21^. For the resilience of urban wildlife it is therefore important that authentic indigenous flora can adapt to urban pressures.

Successful adaptation to the UHIE, as we show here, may also have broader implications for adaptations to climate change. Continued global warming and increased extreme weather, such as intensified heat waves, are predicted for the coming decades^52^, but are already normalised conditions in urban environments^14,15^. The UHIE might therefore act as a selective pressure towards pre-adaptation to climate change^53^. It would be interesting to investigate whether so-called warm equatorial range edge plant populations have a larger fitness in urban heat islands as compared with all other populations^54^. Facilitating UHIE adaptation of naturally occurring flora, for instance via increased connectivity between natural urban and rural areas, could therefore have a positive effect not only on UHIE mitigation in cities but potentially also on plant adaptation to climate change.

## [Author contributions]

YW, KV, BG and KJFV conceptualised the study and KJFV designed the experiments. KJFV collected the field samples. YW and BG collected metadata on the transect locations leading to the subhabitat classification. SI, RK and AJ grew the plants and conducted the experiments. YW and KJFV analysed the data. YW produced the figures and manuscript with input from all authors.

## [Acknowledgements]

We express our appreciation for the work done by members of the GLUE project and, in particular, Jacintha Ellers (VU Amsterdam) who designed the GLUE transect in Amsterdam, on which we based our sampling for this study. We would like to thank Gregor Disveld (NIOO-KNAW) for his assistance in the germination and propagation of plants for greenhouse experiments. Additionally, we appreciate the assistance from Paola Rallo, Cristian Pena and Morgane van Antro (NIOO-KNAW) in watering and scoring of plants in the flowering time experiments. YW is grateful to FLORON for engagement and discussion of the results at their annual symposium, where this work was first presented. This research was funded by the Koninklijke Nederlandse Akademie der Wetenschappen under an institutional grant to the Netherlands Institute of Ecology in collaboration with Naturalis Biodiversity Center.

